# Modelling the impact of urban and hospital wastewaters eco-exposomes on the antibiotic-resistance dynamics

**DOI:** 10.1101/2021.12.17.473112

**Authors:** Paul Henriot, Elena Buelow, Fabienne Petit, Marie-Cécile Ploy, Christophe Dagot, Lulla Opatowski

## Abstract

Antibiotic-resistance emergence and selection have become major public health issues globally. The presence of antibiotic resistant bacteria (ARB) in natural and anthroposophical environments threatens to compromise the sustainability of care in human and animal populations. This study was undertaken to develop a simple model formalizing the selective impact of antibiotics and pollutants on the dynamics of bacterial resistance in water and use the model to analyze longitudinal spatiotemporal data collected in hospital and urban wastewaters. Longitudinal-sampling data were collected between 2012 and 2015 in four different locations in Haute-Savoie, France: hospital and urban wastewaters, before and after water-treatment plants. Concentration in three different types of compounds: 1) heavy metals 2) antibiotics and 3) surfactants; and abundance of 88 individual genes and mobile genetic elements, mostly conferring resistance to antibiotics, were simultaneously collected. A simple hypothesis-driven model describing the weekly ARB dynamics was proposed to fit available data by assuming normalized gene abundance to be proportional to ARB populations in water. Compounds impacts on the dynamics of 17 genes found in multiple sites were estimated. We found that while mercury and vancomycin had relevant effects on ARB dynamics, respectively positively affecting the dynamics of 10 and 12 identified genes, surfactants antagonistically affected genes dynamics (identified for three genes). This simple model enables analyzing the relationship between resistance-gene persistence in aquatic environments and specific compounds inherent to human activities. Applying our model to longitudinal data, we identified compounds that act as co-selectors for antibiotic resistance.

**Highlights:**

- We analyzed longitudinal wastewater resistance genes and environmental data
- We developed a simple hypothesis-driven model to assess resistance selection
- Mercury and vancomycin were key drivers of antibiotic resistance in wastewater

## 1. Introduction

Over the past few decades, the emergence and selection of antibiotic resistance worldwide have become major public health issues. Detection of clinically relevant antibiotic resistance in the environmental bacterial reservoir raises the threat of antimicrobial resistance and the risk of new emergence of antibiotic-resistant pathogens that can colonize humans and animals (Cantas et al., 2013). According to the one health concept, the use of antimicrobial agents in humans, animals and agriculture influences the emergence and selection of antibiotic resistance (Davies and Davies, 2010; Manyi-Loh et al., 2018; Herando-Amado et al., 2019). In the environment, selection of resistant bacteria could be favored by the concomitant presence of bacteria and antibiotics but also could be heightened by co-selection phenomena enhanced by the presence of other chemicals (heavy metals, surfactants, pesticides) (Baker-Austin et al., 2006; Seiler et al., 2012; Devarajan et al., 2015; Singer, et al., 2016; Henriques et al., 2016; Li et al., 2017; Dickinson et al., 2019; Søgaard Jørgensen et al., 2020).

Reflecting human demographics and its related anthropic activities, hospital and urban wastewaters represent major sources of antibiotic-resistant bacteria (ARB), antibiotic-resistance genes (ARGs), and chemical pollutants that contaminate the aquatic environment, despite wastewater treatments (Pärnänen et al., 2019). To date, current understanding of the mechanisms underlying resistance emergence and spread is mainly supported by laboratory experiments (Baharoglu and Mazel, 2011; Gullberg et al., 2014). Identifying the cumulative effects of multiple exposures to contaminants, and assessing their role in the emergence of resistance in natural environments remains a real challenge.

One of the essential inputs to address those combined impacts is to acquire longitudinal data collected at representative sites of anthropized environments that enable analysis through dynamic and mechanistic approaches. In this context, mathematical models formalizing the selection dynamics of resistant bacteria, resistance genes and mobile genetic elements (MGEs)—exposed to compounds—can be devised and used to help quantify and predict selection dynamics in aquatic environments: from wastewater treatment plant (WWTP) effluents to rivers. However, only very few models have been developed to assess driving parameters for the origin of resistance selection in effluents. Existing studies mostly described complex models, taking into account multiple biological or physical mechanisms, were limited by their lack of application to field data (Novozhilov et al., 2005; Ibargüen-Mondragón et al., 2014) or their focus on a single compound (Hellweger et al., 2013; Gothwal and Thatikonda, 2018).

Herein, we used a large spatiotemporal study investigating the presence of chemicals and genetic elements in effluent and affluent hospital and urban wastewaters to develop a simple hypothesis-driven model of antibiotic-resistance selection in anthropized water systems.

## 2. Materials and methods

### 2.1. Data collection and presentation

#### 2.1.1. Study sites

Between 2012 and 2015, data on hospital and urban wastewaters, before and after water treatment, were collected from a national study site in the French Alps (Haute-Savoie). The site, located in Scientrier, France – (46° 06’ 50” North, 6° 19’ 58” East), includes a WWTP with a capacity of 32,000 population equivalents (classical activated sludge system). One biological basin (5400 m^3^) is dedicated to the biological treatment of effluents from the Alpes Leman Hospital Center, which has 450 beds, 8 operating rooms and various medical departments.

Samples were collected in four distinct locations, upstream and downstream (i.e., untreated and treated) of each WWTP (**Fig. 1**) at monthly intervals by flow-proportional sampling. A database with >50,000 entries houses all analytical data (Bertrand-Krajewski et al., 2021). The four sample sites yielded 30, 28, 31 and 19 available time points corresponding to, respectively: untreated and treated hospital wastewaters, and untreated and treated urban wastewaters.

**Fig. 1.**
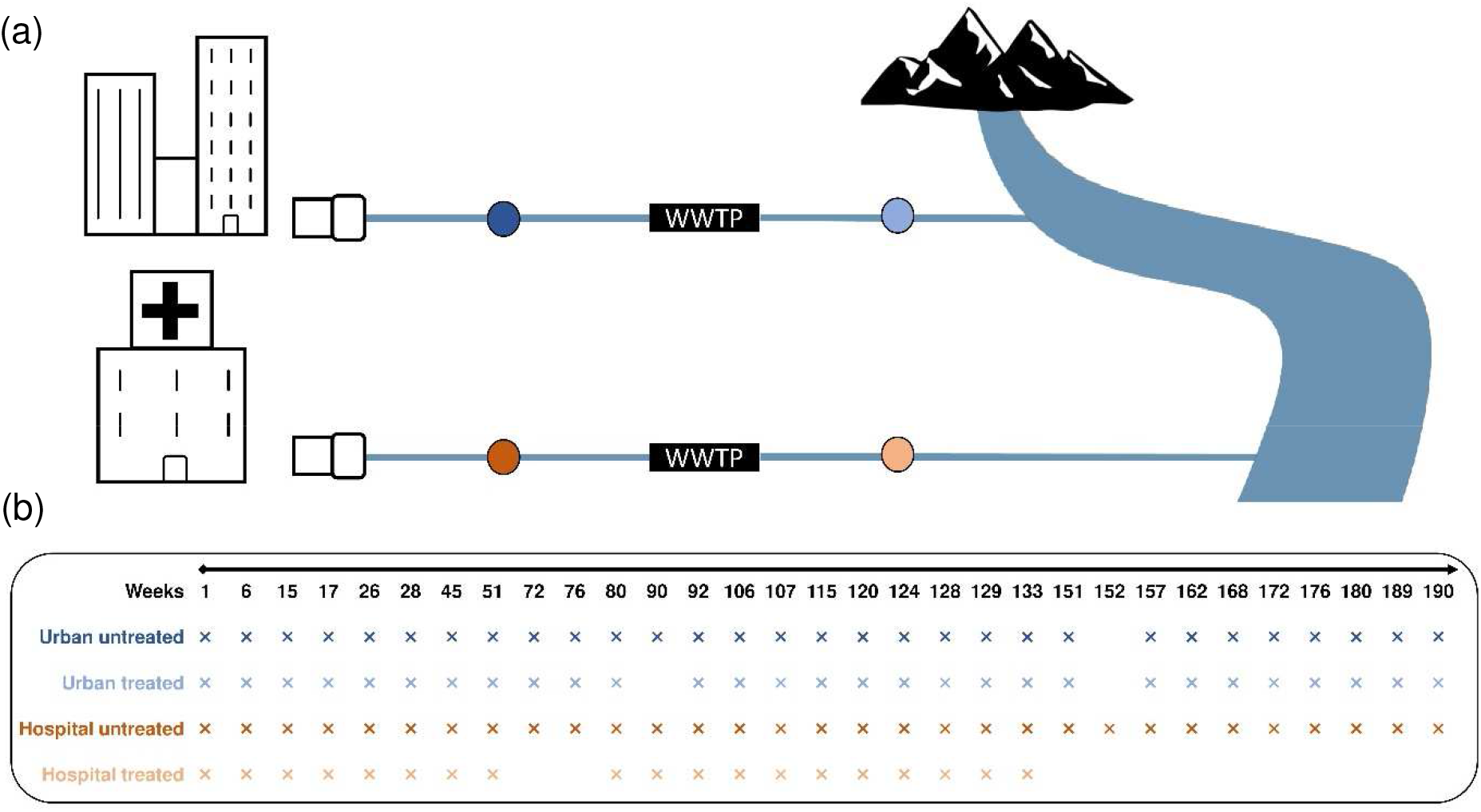
(a) Location of sampling sites (colored circles): the data used were collected between March 2012 and November 2015. This figure is adapted from Buelow et al. (2020). (b) Site-dependent data (88 individual genes and mobile genetic elements, most of them conferring resistance to antibiotics, and some conferring resistance to heavy metals or ammonium compounds) available weekly throughout the sampling period; any missing point corresponds to no available data for that week. WWTP: Wastewater treatment plant

#### 2.1.2. Chemicals

Three different exposure variables were collected simultaneously: (1) heavy metals (ng/L): zinc, nickel, mercury, lead, copper, chromium and cadmium, measured with inductively coupled plasma and atomic emission spectroscopy; (2) antibiotics (ng/L): vancomycin, sulfamethoxazole and ciprofloxacin, measured by solid-phase extraction and liquid chromatography with a tandem mass spectrometry (LC-MS/MS); (3) surfactants (ng/L): anionic, non-ionic and cationic, measured as described by Chonova et al. (2018).

#### 2.1.3. Microbiology

Microorganisms were recovered from filters after water filtration on 45 μm filters and DNA was isolated using the Power Water DNA Extraction Kit (MoBio Laboratories Inc., Carlsbad, CA, USA). Nanoliter-scale quantitative polymerase chain reactions (qPCRs) quantified levels of 88 individually targeted resistance genes and MGEs conferring resistance to antibiotics, surfactants, and/or heavy metals, as described previously (Buelow et al., 2018). qPCR-determined 16S rRNA gene-copy numbers served as a surrogate marker of bacterial biomass, as described previously by Stalder et. al. (2014).

For each sampling time, normalized abundance of 88 individual genes and MGEs, most of them conferring resistance to antibiotics and some conferring resistance to heavy metals or ammonium compounds, were determined (Fig. 1(b)). ARGs, surfactants, heavy metal resistance genes, and MGEs were detected by high-throughput qPCRs, as described above. Individual gene abundance was normalized to the 16S rRNA gene. In our analysis, all values below the limit of quantification (LoQ) were replaced by 0.

More information on sampling, study design, genetic characterization, and chemical analysis are available in Buelow et al. (2020), site descriptions are available in Chonova et al. (2018).

### 2.2. Model choice and hypothesis

#### 2.2.1. Model

We developed a dynamic model to formalize the persistence of resistance genes and MGEs in effluents and the factors affecting their temporal dynamics. We assumed that resistance genes and MGEs were directly associated with the presence of bacteria and, to simplify formalization, modeled the population dynamics of these microbes carrying these genetic elements (**Fig. 2**). In addition, MGEs were considered to be drivers of antibiotic resistance.

**Figure 2.**
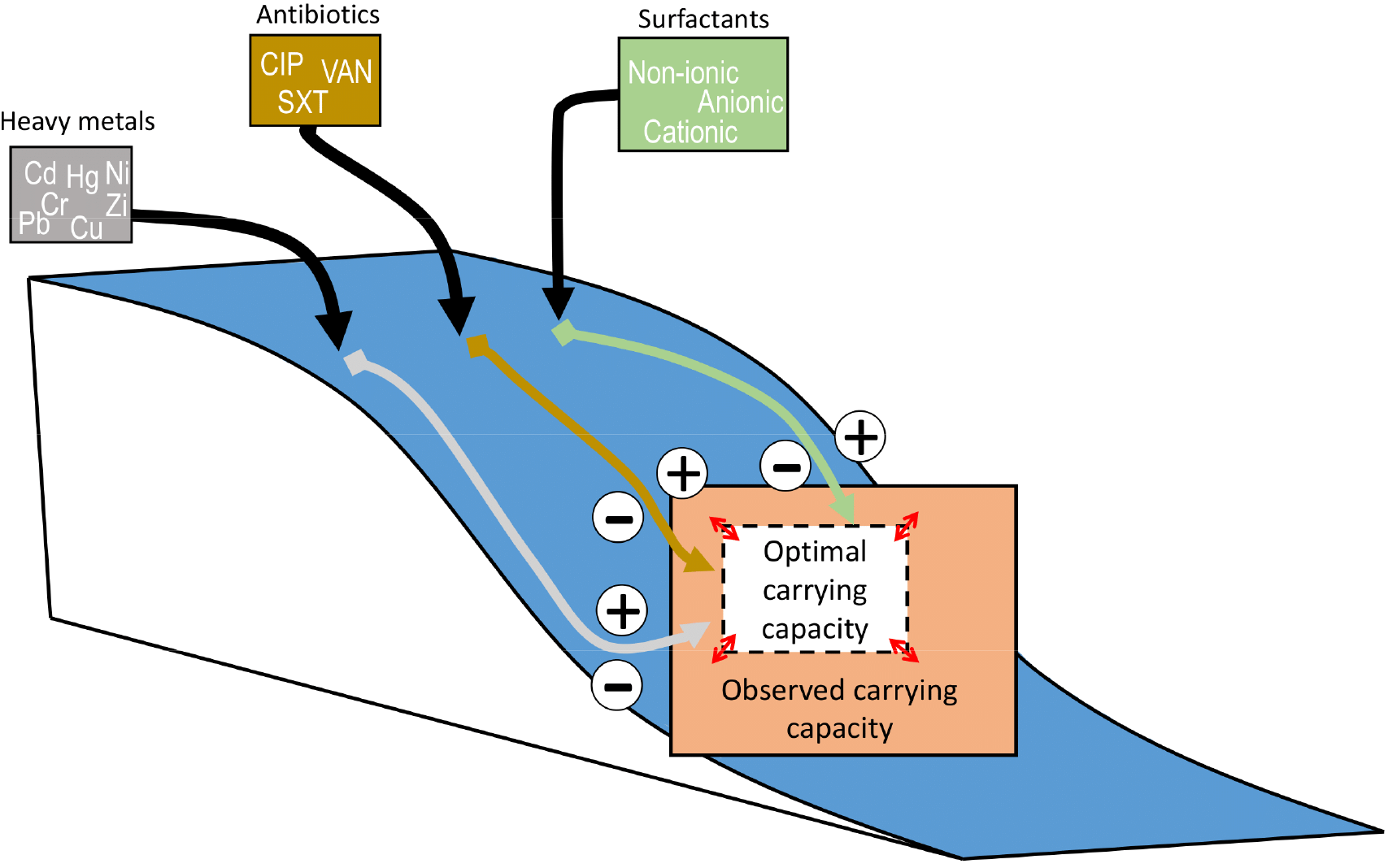
Schematic view of the mechanisms considered in the model. Each compound, varying over time, may have either a selective effect (+) or a reductive effect (–) on resistance status. They directly impact the optimal carrying capacity (*K_opt_*), in the absence of these compounds.

#### 2.2.2. Bacterial population dynamics

We assumed that the dynamic system reached saturated equilibrium every week. We also assumed that the input of resistant bacteria was exogenous and that the available substrate was limited compared to the bacterial biomass, so that almost no bacterial growth occurred and the modeling of bacterial division could be neglected. In addition, we hypothesized that the proportion of free DNA within the sites was negligible and thus did not interfere with the model. Therefore, every week the resistant bacterial population was converted to a stationary state (i.e., *dR/dt* = 0, corresponding to the fixed-point *R*(*t*) = *K(t)*, with *K*(*t*) corresponding to the carrying capacity at time *t* and *R*(*t*) the number of resistant bacteria at time *t*). Hence, the carrying capacity of resistant organisms on a given week *t* was defined as follows:

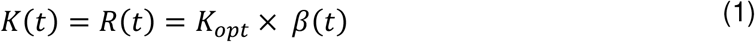

Where *β*(*t*) corresponds to a rate of interaction with the environment at time *t* and *K_opt_* is the optimal carrying capacity in absence of heavy metals, antibiotics and surfactants.

#### 2.2.3. Modeling the impact of compound exposure

Because compound concentrations were measured at different day each week, we assumed that their concentrations were constant for each week, or underwent only small variations that had no impact on the population’s equilibrium level.

We assumed that compounds could have either a selective impact on resistance, thereby increasing the carrying capacity of resistant bacteria, or a reductive impact, thereby reducing the carrying capacity of resistant bacteria.

According to the first hypothesis (called resistance selection), the impact of a given compound, *Cpd*, on *K*(*t*) is written as follows:

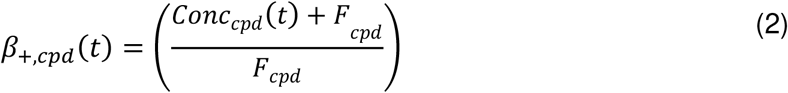

Where *Conc_cpd_*(*t*) is a measured compound concentration at time *t* and *F_cpd_* is a constant selection-threshold concentration value associated with that same agent. When *F_cpd_* is low compared to *Conc_cpd_*(*t*), the selective pressure is high.

According to the second hypothesis (called resistance reduction), the impact of the compound *cpd* on *K*(*t*) is written as follows:

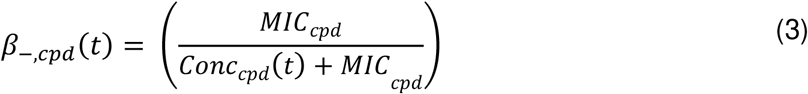

Where *MIC_cpd_* is a constant resistance-reduction threshold concentration value (i.e., minimum inhibitory concentration (MIC)) associated with that same agent. When *MIC_cpd_* is low compared to *Conc_cpd_*(*t*), resistance reduction is high.

Because very little is known about the effect of these different compounds on resistance, we assumed that both processes were possible for all types of agents: antibiotics, heavy metals, and/or surfactants. In the model, including all mechanisms, henceforth called “full”, *β(t)*, including all the possible compounds and their effects in one direction or the other, is expressed as follows:

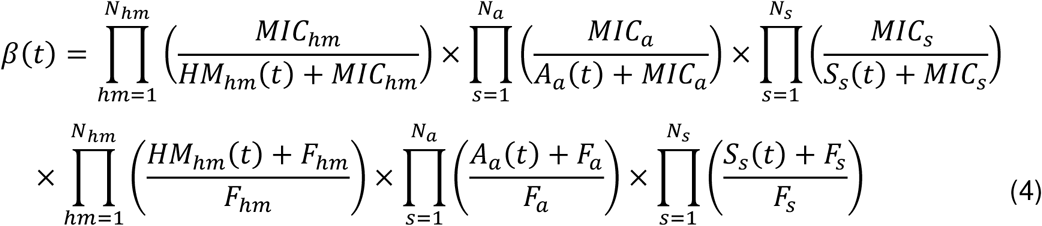

Where resistance-reduction constants are *MIC_hm_* (MIC associated with heavy metal m), *MIC_a_* (MIC associated with antibiotic *a*), and *MIC_S_* (MIC associated with surfactants *s*); selection constants are: *F_hm_* (selection-facilitation constant associated with heavy metal *hm*), *F_a_* (selection-facilitation constant associated with antibiotic *a*), and F_S_(selection-facilitation constant associated with surfactant *s*); *A_a_(t), HM_hm_(t)* and *S_S_(t)* correspond, respectively, to model input variables: antibiotic *a* concentration, heavy metal *hm* exposure concentration and surfactant *s* concentration in the studied environment during week *t*.

Because the full model is complex, with 27 covariables, two sub-models, each incorporating only one of the two mechanisms, were defined and assessed.

Sub-model 1 (i.e., resistance selection), which includes only the amplification effect for all compounds, is written as follows:

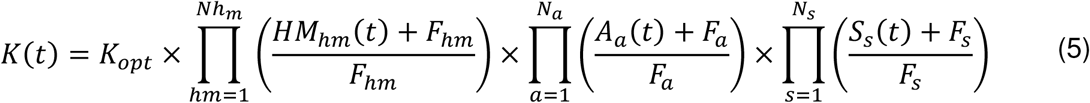

Sub-model 2 (i.e., resistance reduction), which includes only the reductive effect for all compounds, is written as follows:

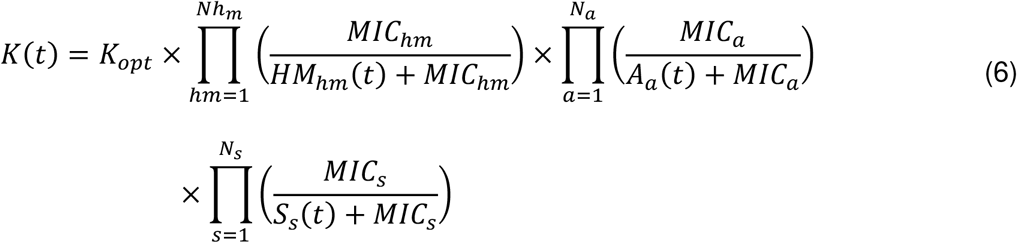

### 2.3. Statistical analyses

Modeling was used to analyze the longitudinal data collected in the four sampling sites. Compound—antibiotic, heavy metal and surfactant—quantities, listed in **Table S1**, were considered model inputs. The quantity of resistant bacteria in the environment at each time, was represented by normalized abundance of genes and MGEs that we considered proportional to the number of resistant bacteria in the environment.

The general model and two sub-models were fitted to the gene- and MGE-abundance data using a Monte-Carlo Markov Chain approach with an adaptive Gibbs sampler. The analysis was run with JAGS software using the R (ver. 4.0.0) interface, “*rjags*”. For each estimation, 300,000 iterations were computed (including 150,000 iterations for the adaptive phase). Chain convergence was assessed through calculation of the potential scale-reduction factor (PSRF; Brooks & Gelman, 1998). The PSRF, noted 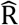, estimates the potential decrease of between-chain variability with respect to the within-chain variability. Convergence was admitted when 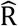 was ≤1.2. For all parameters, non-informative, uniform prior distributions were assumed (see details in **Table A1**), because of lacking *a priori* information on possible parameter values.

In the first step, each of the three models was independently fitted to each of the four site datasets for each resistance gene and MGE. All the model parameters were then estimated for each gene and MGE and each site separately, i.e., 1056 (88 genes × 4 locations × 3 models) independent analyses.

#### 2.3.1. Individual fits

When fitting the model to a specific gene’s dynamics at a given site, it was assessed based on a Gaussian likelihood. Total parameter numbers were 27 for the full model, 14 for sub-models 1 and 2. The likelihood can be written as:

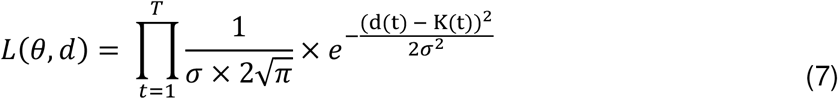

Where *θ* corresponds to the set of parameters, *d*(*t*) is the resistant-bacteria concentration observed at time *t*, *K*(*t*) is the simulated carrying capacity at time *t* (output of the model) and *σ* is the standard deviation, unknown and also estimated.

#### 2.3.2. Model comparisons

For each individual gene, the three models of varying complexity were fitted individually to each site. Models were compared based on a quantitative criterion, the deviance information criterion (*DIC*), which was calculated for each model. The *DIC* enables discriminating the best model by favoring models with good likelihoods, while penalizing the most complex ones (i.e., with more parameters). The *DIC* is calculated as follows:

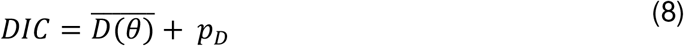

Where *D*(*θ*) = −2 × *log* (*p*(*y*|*θ*)) is the deviance and *p_D_* the effective number of parameters.

The model with the lowest *DIC* is preferred. When two *DIC* values were close (i.e., separated by <5 points), the model with the lower number of parameters was preferred.

#### 2.3.3. Model gene selection

The model fitting was applied to each of the MGEs and the 88 genes encoding resistance to the different types of compounds (heavy metals, antibiotics, surfactants) considered.

For all 1056 individual fits, the following procedure was used to identify the genes with relevant results and discarded all others. For each individual gene fit, the result was considered meaningful when at least two parameters (the optimal carrying capacity, *K_opt_*, plus at least one additional compound-associated constant) were estimated, using prior-posterior distribution overlap (Gimenez et al., 2009) to determine a similarity index between prior and posterior distributions for each estimated parameter. We considered enough information was available in the data to identify a parameter when the distribution overlap was <50%.

In addition, for a given gene, if >30% of data points were below the LoQ (i.e., equal to 0) the corresponding model was excluded.

#### 2.3.4. Assessment of compound impact

To be able to compare estimates across genes within sites, we defined a variable’s strength. For each variable found to have a meaningful positive impact on resistant bacteria selection (i.e., leading to resistance selection), we defined the maximum rate of positive interaction using the 95th percentile value of the compound’s distribution over time, *Cpd*_95%_, and the value of the positive effect of a given compound, *F_cpd_*.

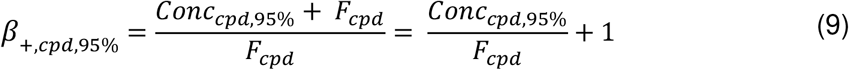

Where *F_cpd_* > 0, therefore *β*_+*cpd*,9K%_ ∈ [1, +∞[. A value close to 1 indicates low positive impact, while a value far from 1 indicates high positive impact.

Similarly, if a variable was found to have a negative impact on resistant-bacteria persistence (i.e., resistance reduction) the maximum negative-interaction rate was calculated as:

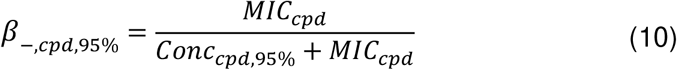

Where *MIC_cpd_* is the value of the negative effect of a given compound, and *Cpd*_95%_ is the 95th percentile of the pollutant-concentration distribution over time and where positive *MIC_cpd_* > 0, therefore *β*_−,*cpd*,95%_ ∈ [0,1]. A value close to 0 indicates high negative impact, while a value close to 1 indicates low negative impact.

## 3. Results

### 3.1. Model selection

The models were used to analyze the observed dynamics for each gene and MGE at each site independently. Resulting *DICs* for each fit and model comparison are summarized in **Table 1** that includes only those genes for which a relevant association was found: only 17 site-specific genes, among a total of 352 analyzed (88 unique genes × 4 sites). Sub-model 1 was selected for 11 of the 17 site-specific genes. The full model was retained for only 5 of the 17 genes. Notably, sub-model 2, which assumes a negative impact of compounds on resistance (i.e., resistance-reduction) was only retained for *tetQ* gene in untreated urban wastewater.

**Table 1.** Individual fits, selected genes and sample sites, and their associated *DICs* obtained by fitting each of the three models.

| Gene | H or U<br>site | Points,<br>$n$ | $DIC$ model | | | Variables<br>involved<br>in dynamics<br>without<br>$K_{opt}$, $n$ |
| --- | --- | --- | --- | --- | --- | --- |
|  |  |  | Full | Sub1: resistance<br>selection | Sub2:<br>resistance<br>reduction |  |
| *aac(3)-II* | U untreated | 30 | **114.03** | 124.99 | 128.57 | 3 |
| *bacA1* | U untreated | 30 | **111.22** | 145.86 | 140.21 | 3 |
| *tetQ* | U untreated | 30 | 282.92 | 285.84 | **279.79** | 1 |
| *aadA* | H untreated | 31 | 295.05 | **293.99** | 317.32 | 1 |
| *aph(3')-Ia, -Ic* | H untreated | 31 | 343.27 | **342.85** | 386.51 | 2 |
| *blaTEM* | H untreated | 31 | **396.22** | 418.15 | 439.72 | 2 |
| *vanB* | H untreated | 31 | 39.66 | **38.29** | 51.73 | 2 |
| *intI2* | H untreated | 31 | 339.06 | **338.08** | 375.19 | 2 |
| *aadE-like* | H treated | 19 | 90.15 | **89.84** | 159.77 | 2 |
| *aph(3')-III* | H treated | 19 | 79.03 | **79.41** | 133.63 | 2 |
| *bacA1* | H treated | 19 | 57.8 | **53.18** | 109.69 | 2 |
| *cblA* | H treated | 19 | **56.41** | 65.23 | 95.39 | 1 |
| *dfrF* | H treated | 19 | 62.33 | **56.9** | 120.2 | 2 |
| *mfsA* | H treated | 19 | 121.63 | **120.47** | 192.27 | 2 |
| *tetO* | H treated | 19 | 121.23 | **121.07** | 182.95 | 2 |
| *tetQ* | H treated | 19 | 146.54 | **139.27** | 188.91 | 2 |
| *tetW* | H treated | 19 | **103.76** | 119.03 | 197.04 | 2 |

H, hospital; U, urban.
For each gene and each site, the selected model is the one with the lowest *DIC* (in bold). When several models yielded similar *DICs*, the simplest one was selected.

Estimated compound effects are provided in Table A2 and estimation results obtained for each of the 17 site-specific genes are summarized in Fig. A1, A2 and A3. Among all variables entered into our model, mercury was relevantly associated with 10 out of 17 genes and MGEs; suggesting a selection-facilitation effect on the bacterial resistance level, mostly in Hospital treated water; vancomycin was the most frequently relevant antibiotic, selected for 13 out of 17 genes and MGEs; and anionic surfactants were the most impacting among surfactants (significant for 3 out of 17 genes). Our modeling results suggested that mercury and vancomycin acted as resistance-selection facilitators, while anionic surfactant reduced resistance selection.

Effects of those three compounds for the different genes and sites are summarized in Fig. 3a-c, together with the fitted dynamics of three relevant selected resistance-gene associated with them: *bacA1* in treated wastewater, *blaTEM* in hospital untreated wastewater, and *aac3* in urban untreated wastewater (Fig. 3d-f).

**Figure 3.**
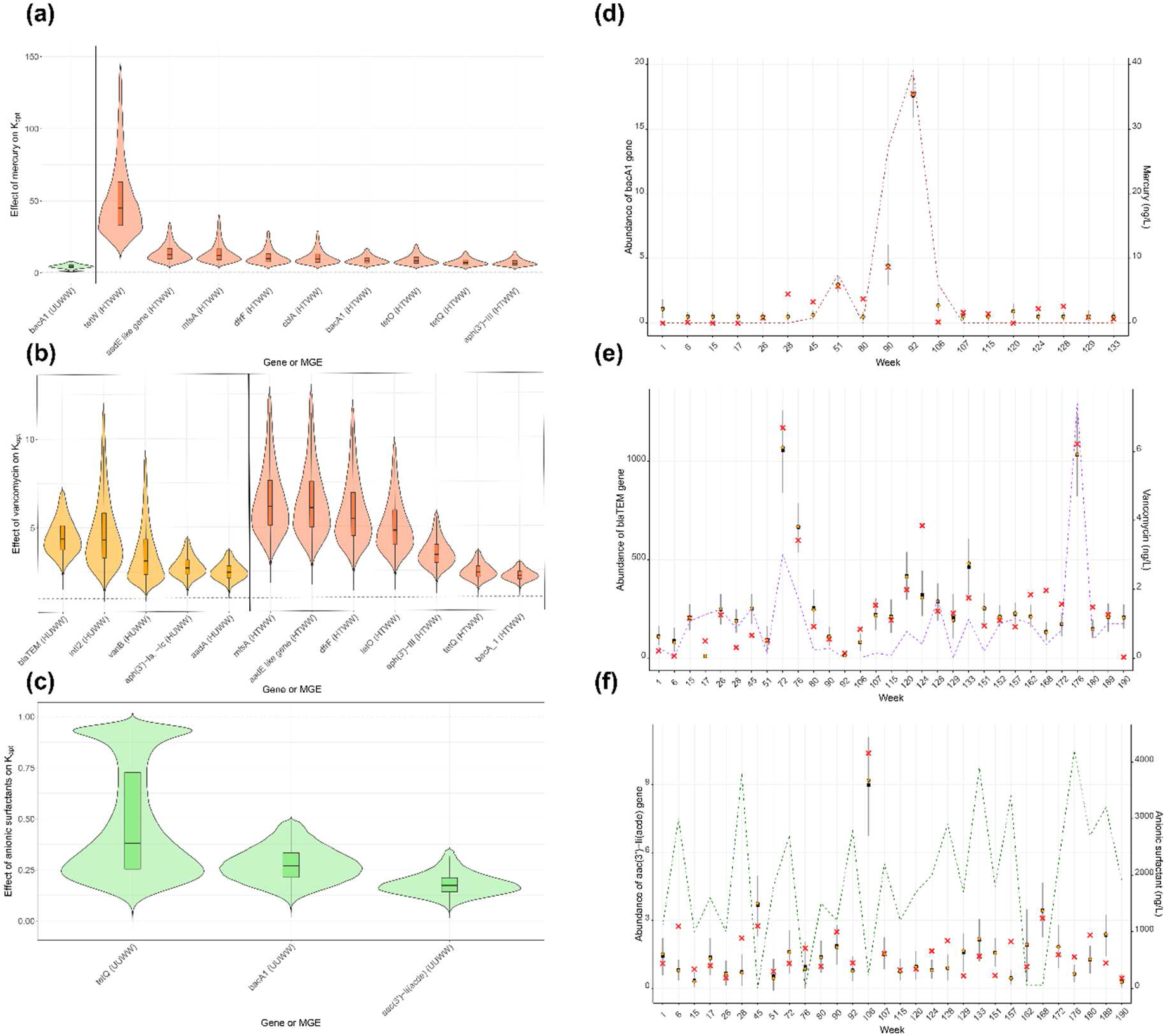
Model fits to the collected data for resistance dynamics of selected genes and MGEs and compound concentrations. (a-c) Normalized plots of the effect*s* of selected compounds on *K_opt_* (optimal carrying capacity) as a reflection of the estimated strength associated with a given agent for all genes and sites: (a) mercury, (b) vancomycin and (c) anionic surfactant. The box plots report the median (internal horizontal line), 25th and 75th percentiles (lower and upper limits) and the 2.5th and 97.5th distribution percentiles (vertical lines). (d–f) Predicted (in the best-performing model) and reported relative resistance-gene and MGE abundance (left *y*-axis) and selected associated compound concentration (right *y*-axis). Red crosses indicate collected data, black squares the median prediction, blue vertical lines the 95% prediction intervals, and orange circles the *K*-distribution modes at each time *t*. (d) *bacA1* over time in treated hospital wastewater, with trends (dashed lines) for detected mercury concentrations. (e) *blaTEM* over time in untreated hospital wastewater, with trends for detected vancomycin concentrations. (f) *aac3* in untreated urban wastewater, with trends for detected anionic surfactant concentrations.

### 3.2. Mercury

For the *bacA1*-gene, the estimated median interaction rate for mercury was 8.67 (Fig. 3a), meaning that when mercury reaches its 95% highest measured concentration of 28.2 ng/L, the population size of bacteria harboring the *bacA1*-gene was increased by a factor of 8.67 (in median). For the other selected genes, the median effect varied widely, from 6.91 (*aph(3)-III gene*) to 46.37 (*tetW gene*) in treated hospital water. Some effect was also detected for *bacA1* gene (median 4.62) in untreated urban wastewater.

### 3.3. Vancomycin

This antibiotic had a meaningful effect on the dynamics of *blaTEM*, a β-lactam resistance gene, based on data collected in untreated hospital wastewater. Predicted dynamics and observed resistance are shown in Fig. 3b. An estimated vancomycin concentration of 2.37 ng/L generates a 4.47-fold increase of the initial carrying capacity of bacteria harboring *blaTEM*. The median vancomycin effect ranged from 2.47 (*aadA gene*) to 4.47 (*blaTEM gene*) in untreated hospital water, where relevant effects were also detected for the following genes and MGEs: *aph(3’)-Ia -Ic, vanB and IntI2. In the hospital treated wastewater* relevant effects of vancomycin was detected for *bacA1, tetQ, aph(3)-III, tetO, dfrF, mfsA* genes, and an *aadE*-like gene, with median effects ranging from 2.18 to 6.22.

### 3.4. Anionic surfactants

Finally, anionic surfactants inhibited the dynamics of three genes the in untreated urban wastewater (Fig. 3e). At a concentration of 3855 ng/L, an almost 80% reduction (median 0.195) of the initial carrying capacity was observed for bacteria harboring the *aac(3)-II* gene. For all selected genes, the median estimated effect ranged from 0.195 (*aac(3’)-II gene*) to 0.395 (*tetQ gene*).

## 4. Discussion

### 4.1. Main findings

Herein, we proposed a novel modeling framework that simulates the relationships between the persistence of bacterial antibiotic-resistance genes in wastewater before and after a WWTP and the detection of specific compounds inherent in human activities. Using longitudinal data collected within a large temporal analytical study and bringing together determinations of antibiotic-resistance gene presence in different types of wastewaters, and characterizations of environmental exposomes, we estimated the strengths of selective or reductive effects of various compounds on the 88 resistance genes and MGEs evaluated.

First, our results demonstrated that a relatively simple mathematical representation was able to identify compound-interaction signals for 17 site-specific genes and MGEs out of 88 assessed at four sites and 13 chemical agents. Hypothesizing that chemicals present in wastewater could be key drivers of resistance in bacterial population dynamics in such aquatic systems, we were able to identify, from our modeling results, an association between some players of the exposome and the evolutions of resistance-gene abundance.

Mercury and vancomycin provided a selective advantage to a series of resistance genes and MGEs. Identifying such drivers is key to reducing the concentrations of co-selecting molecules, thereby limiting resistance-gene selection in such aquatic environments.

### 4.2. Heavy metals

According to our models, mercury most strongly selected several antibiotic-resistance genes, for instance for multiple genes encoding resistance to tetracycline (*tetW, tetQ* and *tetO* in treated hospital wastewater). That finding is in line with previously reported studies on mercury’s impact on antibiotic-resistance selection (Skurnik et al., 2010). Lloyd et al. (2018) showed that mercury-exposed fish microbiomes in the environment were highly susceptible to containing antibiotic-multiresistant bacteria, compared to unexposed controls. Co-resistance due to mercury is apparently not new, since Wardwell et al. (2009) showed that mercury-exposed indigenous bacterial strains in 2000-year-old sphagnum core samples had a multidrug-resistance phenotype.

Interestingly, chromium and cadmium, respectively, antagonistically affected *bacA1*-gene (in untreated urban wastewater) and *blaTEM-gene* dynamics (in untreated hospital wastewater). Because gene appearance is often dependent on bacterial species, those results could be attributable to specific inhibition of antibiotic-sensitive strains. In addition, some resistance gene carriage could represent a fitness cost that would confer reactivity to some metal compounds.

Our results are consistent with previous findings highlighting the selective impact of heavy metals on resistance. Seiler and Berendonk (2012) conducted a risk assessment to determine potential metal-driven co-selection, and found that mercury, zinc, copper and cadmium were potentially important factors affecting co-selection for antibiotic resistance. The results of another modeling study (Hellweger et al., 2013) suggested that metals could co-select for tetracycline resistance. Those authors also found signals for zinc and copper that were not confirmed herein.

### 4.3. Antibiotics

According to our models, among the three antibiotic concentrations tracked here, vancomycin was found to most closely impact antibiotic resistance. We found a significant association between vancomycin and the temporal dynamics of 14 different site-specific genetic elements. Interestingly those were not exclusively involved in vancomycin resistance found in untreated and treated hospital wastewaters. Those results are in line with previous studies showing strong selection of vancomycin resistance under over-exposure to vancomycin (Cetinkaya et al., 2000; Hiramatsu, 2001). In addition, this antibiotic could act as a co-selector for other resistance genes, assuming that some vancomycin-resistance genes might be part of large transferable genetic elements (Poyart et al., 1997; Bender et al., 2016; Ahmed and Baptiste, 2018).

The fact that our model predicts vancomycin to also act as a facilitator for non-vancomycin-resistance genes (mostly carried by Gram-positive bacteria) may have other sources of explanation. First, vancomycin could have led to inhibition of Gram-positive bacteria susceptible to this antibiotic and allowed an increase in Gram-negative bacteria, naturally resistant to vancomycin and carrying other antibiotic resistance genes (for example blaTEM). Second, simultaneous in-hospital use of other antibiotics might have conducted to their release into the water but they were not measured or not detected because they are generally less persistent in the environment.

Thus, these non-vancomycin-resistance–gene abundance might be concomitant with vancomycin concentrations; associations between vancomycin and some resistance genes would rather result from selective pressure of those undetected antibiotics. Here, vancomycin would be playing a role in proxy antibiotic contamination in the hospital wastewater environments analyzed.

### 4.4. Surfactants

Interestingly, model results associated anionic and non-ionic surfactants with resistance reduction for some genes (i.e., decreasing specific resistance gene abundance) whereas no association was found for cationic surfactants. Anionic compounds were reductive of certain resistance genes, like *bacA1* (in untreated urban wastewater), where non-ionic surfactants also had a restrictive effect, or *tetQ* (in untreated urban wastewater). The involved genes are mostly carried by Gram-negative bacteria, like *tetQ* or *aac3* (Leng et al., 1997; Nie et al., 2014; Lalitha Aishwarya et al., 2020; Jasemi, et al. 2021). Therefore, these resistance genes might reflect the presence of Gram-negative bacteria in urban untreated wastewater, suggesting that surfactants might induce cell death in these bacteria.

### 4.5. Limitations

Several modeling assumptions and limitations of our study warrant discussion. First, because the model was specifically developed to adapt to the available data, it is rather limited by its simplicity. Bacterial growth was taken to be negligible and we only analyzed the stationary weekly state. In the future, consideration of the within-week variation of the eco-exposome could potentially improve the model and provide more accurate results.

Second, despite assessment of three integrase-encoding genes and nine transposase genes associated with high gene transfer and antibiotic resistance, information regarding the bacteria (i.e., genus and species) were not available. That lack prevented specific reproduction of some important transmission mechanisms, like horizontal and vertical transfer or transposition.

Third, our analysis included only a limited number of compounds, and did not consider temperature, water flows or other variables (i.e., local epidemiology or antibiotic prescriptions). Future studies should include more variables to avoid possible confounding effects.

Finally, for each independent model fit, relevant effects were selected using the prior-posterior overlap method, with a 50% threshold. Because that coverage was rather arbitrary, a 35% threshold was tested, as proposed in Gimenez et al. (2009); the results were unchanged.

Our results suggest that water composition highly depends on the sampling site, where compounds are strongly specific. Indeed, no genetic element could be identified as a general indicator of antibiotic-resistance in the anthropized wastewaters analyzed, even considering hospital and urban wastewaters separately.

## 5. Conclusion

Herein, we described a new mechanistic modeling framework that helps better understand of the role of various compounds in antibiotic-resistance selection in bacteria in aquatic environments. Our results suggest co-selection for antibiotic resistance by the eco-exposome, specifically by mercury and vancomycin.

## Abbreviations

ARB: antibiotic resistant bacteria
ARG: antibiotic resistance gene
MGE: mobile genetic element
WWTP: wastewater treatment plant
PSRF: potential scale-reduction factor

## Contribution

**PH**, **LO** and **CD** designed the study. **CD** provided access to data. **EB** provided support for data description and interpretation. **PH** built the model and did the analyses with input from **LO**. **PH**, **EB**, **LO** and **CD** wrote the manuscript with contribution of **FP** and **MCP**.

## Declaration of competing interest

We declare no competing interests.

## Acknowledgments

We would like to thank the SIPIBEL consortium for displaying data.

## Funding

This study was funded by Agence Nationale Sécurité Sanitaire Alimentaire Nationale (Pandore Project, EST 2017/3 ABR/28). It was also supported directly by internal resources from the French National Institute for Health and Medical Research, the Institut Pasteur, and the University of Versailles–Saint-Quentin-en-Yvelines / University of Paris-Saclay. This study received funding from the French Government’s ‘Investissement d’Avenir’ program, Laboratoire d’Excellence ‘Integrative Biology of Emerging Infectious Diseases’ (Grant ANR-10-LABX-62-IBEID).

## Notes

### Competing Interest Statement

The authors have declared no competing interest.

## References

Ahmed, M.O., Baptiste, K.E., 2018. Vancomycin-Resistant Enterococci: A Review of Antimicrobial Resistance Mechanisms and Perspectives of Human and Animal Health. Microbial Drug Resistance 24, 590–606. https://doi.org/10.1089/mdr.2017.0147

Baharoglu, Z., Mazel, D., 2011. Vibrio cholerae Triggers SOS and Mutagenesis in Response to a Wide Range of Antibiotics: a Route towards Multiresistance. Antimicrob Agents Chemother 55, 2438–2441. https://doi.org/10.1128/AAC.01549-10

Baker-Austin, C., Wright, M.S., Stepanauskas, R., McArthur, J.V., 2006. Co-selection of antibiotic and metal resistance. Trends in Microbiology 14, 176–182. https://doi.org/10.1016/j.tim.2006.02.006

Bender, J.K., Kalmbach, A., Fleige, C., Klare, I., Fuchs, S., Werner, G., 2016. Population structure and acquisition of the vanB resistance determinant in German clinical isolates of Enterococcus faecium ST192. Sci Rep 6, 21847. https://doi.org/10.1038/srep21847

Bertrand-Krajewski, Jean-Luc, Bournique, Rémy, Lecomte, Vivien, Pernin, Noémie, Wiest, Laure, Bazin, Christine, Bouchez, Agnès, Brelot, Elodie, Cournoyer, Benoît, Chonova, Teofana, Dagot, Christophe, Di Majo, Pascal, Gonzalez-Ospina, Adriana, Klein, Audrey, Labanowski, Jérôme, Levi, Yves, Perodin, Yves, Rabello-Vargas, Sandra, Reuilly, Liana, Rocj, Audrey, Wahl, Alex, 2021. SIPIBEL data set. https://doi.org/10.5281/ZENODO.5176161

Brooks, S.P., Gelman, A., 1998. General Methods for Monitoring Convergence of Iterative Simulations. Journal of Computational and Graphical Statistics 7, 434–455. https://doi.org/10.1080/10618600.1998.10474787

Buelow, E., Bayjanov, J.R., Majoor, E., Willems, R.J., Bonten, M.J., Schmitt, H., van Schaik, W., 2018. Limited influence of hospital wastewater on the microbiome and resistome of wastewater in a community sewerage system. FEMS Microbiology Ecology 94. https://doi.org/10.1093/femsec/fiy087

Buelow, E., Rico, A., Gaschet, M., Lourenço, J., Kennedy, S.P., Wiest, L., Ploy, M.-C., Dagot, C., 2020. Hospital discharges in urban sanitation systems: Long-term monitoring of wastewater resistome and microbiota in relationship to their eco-exposome. Water Research X 7, 100045. https://doi.org/10.1016/j.wroa.2020.100045

Cantas, L., Shah, S.Q.A., Cavaco, L.M., Manaia, C.M., Walsh, F., Popowska, M., Garelick, H., Bürgmann, H., Sørum, H., 2013. A brief multi-disciplinary review on antimicrobial resistance in medicine and its linkage to the global environmental microbiota. Front. Microbiol. 4. https://doi.org/10.3389/fmicb.2013.00096

Cetinkaya, Y., Falk, P., Mayhall, C.G., 2000. Vancomycin-Resistant Enterococci. Clin Microbiol Rev 13, 686–707. https://doi.org/10.1128/CMR.13.4.686

Chonova, T., Lecomte, V., Bertrand-Krajewski, J.-L., Bouchez, A., Labanowski, J., Dagot, C., Lévi, Y., Perrodin, Y., Wiest, L., Gonzalez-Ospina, A., Cournoyer, B., Sebastian, C., 2018. The SIPIBEL project: treatment of hospital and urban wastewater in a conventional urban wastewater treatment plant. Environ Sci Pollut Res 25, 9197–9206. https://doi.org/10.1007/s11356-017-9302-0

Davies, J., Davies, D., 2010. Origins and Evolution of Antibiotic Resistance. Microbiol Mol Biol Rev 74, 417–433. https://doi.org/10.1128/MMBR.00016-10

Devarajan, N., Laffite, A., Graham, N.D., Meijer, M., Prabakar, K., Mubedi, J.I., Elongo, V., Mpiana, P.T., Ibelings, B.W., Wildi, W., Poté, J., 2015. Accumulation of Clinically Relevant Antibiotic-Resistance Genes, Bacterial Load, and Metals in Freshwater Lake Sediments in Central Europe. Environ. Sci. Technol. 49, 6528–6537. https://doi.org/10.1021/acs.est.5b01031

Dickinson, A.W., Power, A., Hansen, M.G., Brandt, K.K., Piliposian, G., Appleby, P., O’Neill, P.A., Jones, R.T., Sierocinski, P., Koskella, B., Vos, M., 2019. Heavy metal pollution and co-selection for antibiotic resistance: A microbial palaeontology approach. Environment International 132, 105117. https://doi.org/10.1016/j.envint.2019.105117

Gimenez, O., Morgan, B.J.T., Brooks, S.P., 2009. Weak Identifiability in Models for Mark-Recapture-Recovery Data, in: Thomson, D.L., Cooch, E.G., Conroy, M.J. (Eds.), Modeling Demographic Processes In Marked Populations. Springer US, Boston, MA, pp. 1055–1067. https://doi.org/10.1007/978-0-387-78151-8_48

Gothwal, R., Thatikonda, S., 2018. Mathematical model for the transport of fluoroquinolone and its resistant bacteria in aquatic environment. Environ Sci Pollut Res 25, 20439–20452. https://doi.org/10.1007/s11356-017-9848-x

Gullberg, E., Albrecht, L.M., Karlsson, C., Sandegren, L., Andersson, D.I., 2014. Selection of a Multidrug Resistance Plasmid by Sublethal Levels of Antibiotics and Heavy Metals. mBio 5. https://doi.org/10.1128/mBio.01918-14

Hellweger, F., Ruan, X., Sanchez, S., 2011. A Simple Model of Tetracycline Antibiotic Resistance in the Aquatic Environment (with Application to the Poudre River). IJERPH 8, 480–497. https://doi.org/10.3390/ijerph8020480

Henriques, I., Tacão, M., Leite, L., Fidalgo, C., Araújo, S., Oliveira, C., Alves, A., 2016. Co-selection of antibiotic and metal(loid) resistance in gram-negative epiphytic bacteria from contaminated salt marshes. Marine Pollution Bulletin 109, 427–434. https://doi.org/10.1016/j.marpolbul.2016.05.031

Hernando-Amado, S., Coque, T.M., Baquero, F., Martínez, J.L., 2019. Defining and combating antibiotic resistance from One Health and Global Health perspectives. Nat Microbiol 4, 1432–1442. https://doi.org/10.1038/s41564-019-0503-9

Hiramatsu, K., 2001. Vancomycin-resistant Staphylococcus aureus: a new model of antibiotic resistance. The Lancet Infectious Diseases 1, 147–155. https://doi.org/10.1016/S1473-3099(01)00091-3

Ibargüen-Mondragón, E., Mosquera, S., Cerón, M., Burbano-Rosero, E.M., Hidalgo-Bonilla, S.P., Esteva, L., Romero-Leitón, J.P., 2014. Mathematical modeling on bacterial resistance to multiple antibiotics caused by spontaneous mutations. Biosystems 117, 60–67. https://doi.org/10.1016/j.biosystems.2014.01.005

Jasemi, S., Emaneini, M., Ahmadinejad, Z., Fazeli, M.S., Sechi, L.A., Sadeghpour Heravi, F., Feizabadi, M.M., 2021. Antibiotic resistance pattern of Bacteroides fragilis isolated from clinical and colorectal specimens. Ann Clin Microbiol Antimicrob 20, 27. https://doi.org/10.1186/s12941-021-00435-w

Lalitha Aishwarya, K.V., Venkataramana Geetha, P., Eswaran, S., Mariappan, S., Sekar, U., 2020. Spectrum of Aminoglycoside Modifying Enzymes in Gram-Negative Bacteria Causing Human Infections. J Lab Physicians 12, 27–31. https://doi.org/10.1055/s-0040-1713687

Leng, Z., Riley, D.E., Berger, R.E., Krieger, J.N., Roberts, M.C., 1997. Distribution and mobility of the tetracycline resistance determinant tetQ. J Antimicrob Chemother 40, 551–559. https://doi.org/10.1093/jac/40.4.551

Li, L.-G., Xia, Y., Zhang, T., 2017. Co-occurrence of antibiotic and metal resistance genes revealed in complete genome collection. ISME J 11, 651–662. https://doi.org/10.1038/ismej.2016.155

Lloyd, N.A., Nazaret, S., Barkay, T., 2018. Whole genome sequences to assess the link between antibiotic and metal resistance in three coastal marine bacteria isolated from the mummichog gastrointestinal tract. Marine Pollution Bulletin 135, 514–520. https://doi.org/10.1016/j.marpolbul.2018.07.051

Manyi-Loh, C., Mamphweli, S., Meyer, E., Okoh, A., 2018. Antibiotic Use in Agriculture and Its Consequential Resistance in Environmental Sources: Potential Public Health Implications. Molecules 23, 795. https://doi.org/10.3390/molecules23040795

Nie, L., Lv, Y., Yuan, M., Hu, X., Nie, T., Yang, X., Li, G., Pang, J., Zhang, J., Li, C., Wang, X., You, X., 2014. Genetic basis of high level aminoglycoside resistance in Acinetobacter baumannii from Beijing, China. Acta Pharmaceutica Sinica B 4, 295–300. https://doi.org/10.1016/j.apsb.2014.06.004

Novozhilov, A.S., 2005. Mathematical Modeling of Evolution of Horizontally Transferred Genes. Molecular Biology and Evolution 22, 1721–1732. https://doi.org/10.1093/molbev/msi167

Pärnänen, K.M.M., Narciso-da-Rocha, C., Kneis, D., Berendonk, T.U., Cacace, D., Do, T.T., Elpers, C., Fatta-Kassinos, D., Henriques, I., Jaeger, T., Karkman, A., Martinez, J.L., Michael, S.G., Michael-Kordatou, I., O’Sullivan, K., Rodriguez-Mozaz, S., Schwartz, T., Sheng, H., Sørum, H., Stedtfeld, R.D., Tiedje, J.M., Giustina, S.V.D., Walsh, F., Vaz-Moreira, I., Virta, M., Manaia, C.M., 2019. Antibiotic resistance in European wastewater treatment plants mirrors the pattern of clinical antibiotic resistance prevalence. Sci. Adv. 5, eaau9124. https://doi.org/10.1126/sciadv.aau9124

Poyart, C., Pierre, C., Quesne, G., Pron, B., Berche, P., Trieu-Cuot, P., 1997. Emergence of vancomycin resistance in the genus Streptococcus: characterization of a vanB transferable determinant in Streptococcus bovis. Antimicrob Agents Chemother 41, 24–29. https://doi.org/10.1128/AAC.41.1.24

Seiler, C., Berendonk, T.U., 2012. Heavy metal driven co-selection of antibiotic resistance in soil and water bodies impacted by agriculture and aquaculture. Front. Microbio. 3. https://doi.org/10.3389/fmicb.2012.00399

Singer, A.C., Shaw, H., Rhodes, V., Hart, A., 2016. Review of Antimicrobial Resistance in the Environment and Its Relevance to Environmental Regulators. Front. Microbiol. 7. https://doi.org/10.3389/fmicb.2016.01728

Skurnik, D., Ruimy, R., Ready, D., Ruppe, E., Bernède-Bauduin, C., Djossou, F., Guillemot, D., Pier, G.B., Andremont, A., 2010. Is exposure to mercury a driving force for the carriage of antibiotic resistance genes? Journal of Medical Microbiology 59, 804–807. https://doi.org/10.1099/jmm.0.017665-0

Søgaard Jørgensen, P., Folke, C., Henriksson, P.J.G., Malmros, K., Troell, M., Zorzet, A., 2020. Coevolutionary Governance of Antibiotic and Pesticide Resistance. Trends in Ecology & Evolution 35, 484–494. https://doi.org/10.1016/j.tree.2020.01.011

Stalder, T., Barraud, O., Jové, T., Casellas, M., Gaschet, M., Dagot, C., Ploy, M.-C., 2014. Quantitative and qualitative impact of hospital effluent on dissemination of the integron pool. ISME J 8, 768–777. https://doi.org/10.1038/ismej.2013.189

Wardwell, L.H., Jude, B.A., Moody, J.P., Olcerst, A.I., Gyure, R.A., Nelson, R.E., Fekete, F.A., 2009. Co-Selection of Mercury and Antibiotic Resistance in Sphagnum Core Samples Dating Back 2000 Years. Geomicrobiology Journal 26, 351–360. https://doi.org/10.1080/01490450902889072

