## Supplementary appendix for "Modelling the impact of urban and hospital wastewaters eco-exposomes on the antibiotic-resistance dynamics"

**Supplementary appendix A**

**Table A.1.** Uniform prior upper and lower limits.

| Parameter | Lower limit | Upper limit |
| --- | --- | --- |
| *K_opt_* | 10^–7^ | 10,000 |
| *A_CIP_* | 10^–7^ | 10,000 |
| *A_SXT_* | 10^–7^ | 1,000 |
| *A_VAN_* | 10^–7^ | 1,000 |
| *HM_Cd_* | 10^–7^ | 1,000 |
| *HM_Cr_* | 10^–7^ | 1,000 |
| *HM_Cu_* | 10^–7^ | 10,000 |
| *HM_Pb_* | 10^–7^ | 1,000 |
| *HM_Hg_* | 10^–7^ | 10,000 |
| *HM_Ni_* | 10^–7^ | 1,000 |
| *HM_Zn_* | 10^–7^ | 100,000 |
| *S_anionnic_* | 10^–7^ | 1,000,000 |
| *S_cationic_* | 10^–7^ | 1,000,000 |
| *S_non-ionic_* | 10^–7^ | 1,000,000 |

K_opt_, optimal carrying capacity; A, antibiotic; HM, heavy metal, S, surfactant;

CIP, ciprofloxacin; SXT, sulfamethoxazole; VAN, vancomycin; Cd, cadmium;

Cr, chromium; Cu copper; Pb, lead, Hg, mercury, Ni, nickel; Zn, zinc

**Table A.2.** Estimated effects of compound-associated parameters for each selected gene with prior-posterior overlap <50% (colored rectangles).

| Gene | H or U site |  | Antibiotic | | |  | Heavy metal | | | | | | | |  | | Surfactant | | |
| --- | --- | --- | --- | --- | --- | --- | --- | --- | --- | --- | --- | --- | --- | --- | --- | --- | --- | --- | --- |
|  |  | Resistance | CIP | SXT | VAN |  | Cd | Cr | Cu | Pb | Hg | Ni | Zi |  | | Anionic | | Cationic | Non-ionic |
| *aac(3')-II* | U untreated | Aminoglycosides |  |  |  |  |  |  |  |  |  |  |  |  | |  | |  |  |
| *bacA 1* | U untreated | Bacitracin |  |  |  |  |  |  |  |  |  |  |  |  | |  | |  |  |
| *tetQ* | U untreated | Tetracycline |  |  |  |  |  |  |  |  |  |  |  |  | |  | |  |  |
| *aadA* | H untreated | Aminoglycosides |  |  |  |  |  |  |  |  |  |  |  |  | |  | |  |  |
| *aph(3')-Ia, -Ic* | H untreated | Aminoglycosides |  |  |  |  |  |  |  |  |  |  |  |  | |  | |  |  |
| *blaTEM* | H untreated | β-lactams |  |  |  |  |  |  |  |  |  |  |  |  | |  | |  |  |
| *vanB* | H untreated | Vancomycin |  |  |  |  |  |  |  |  |  |  |  |  | |  | |  |  |
| *intI2* | H untreated | Multiple |  |  |  |  |  |  |  |  |  |  |  |  | |  | |  |  |
| *aadE-like gene* | H treated | Aminoglycosides |  |  |  |  |  |  |  |  |  |  |  |  | |  | |  |  |
| *aph(3')-III* | H treated | Aminoglycosides |  |  |  |  |  |  |  |  |  |  |  |  | |  | |  |  |
| *bacA 1* | H treated | Bacitracin |  |  |  |  |  |  |  |  |  |  |  |  | |  | |  |  |
| *cblA* | H treated | β-lactams |  |  |  |  |  |  |  |  |  |  |  |  | |  | |  |  |
| *dfrF* | H treated | Trimethoprim |  |  |  |  |  |  |  |  |  |  |  |  | |  | |  |  |
| *mfsA* | H treated | Multiple |  |  |  |  |  |  |  |  |  |  |  |  | |  | |  |  |
| *tetO* | H treated | Tetracycline |  |  |  |  |  |  |  |  |  |  |  |  | |  | |  |  |
| *tetQ* | H treated | Tetracycline |  |  |  |  |  |  |  |  |  |  |  |  | |  | |  |  |
| *tetW* | H treated | Tetracycline |  |  |  |  |  |  |  |  |  |  |  |  | |  | |  |  |

Blue, resistance-elimination effect; red, resistance-selection effect. If overlap exceeded 50%, the parameter was considered to be weakly identifiable

regarding the available data (grey rectangles). H, hospital; U, urban, CIP, ciprofloxacin; SXT, sulfamethoxazole; VAN, vancomycin; Cd, cadmium; Cr, chromium;

Cu copper; Pb, lead, Hg, mercury, Ni, nickel; Zn, zinc.

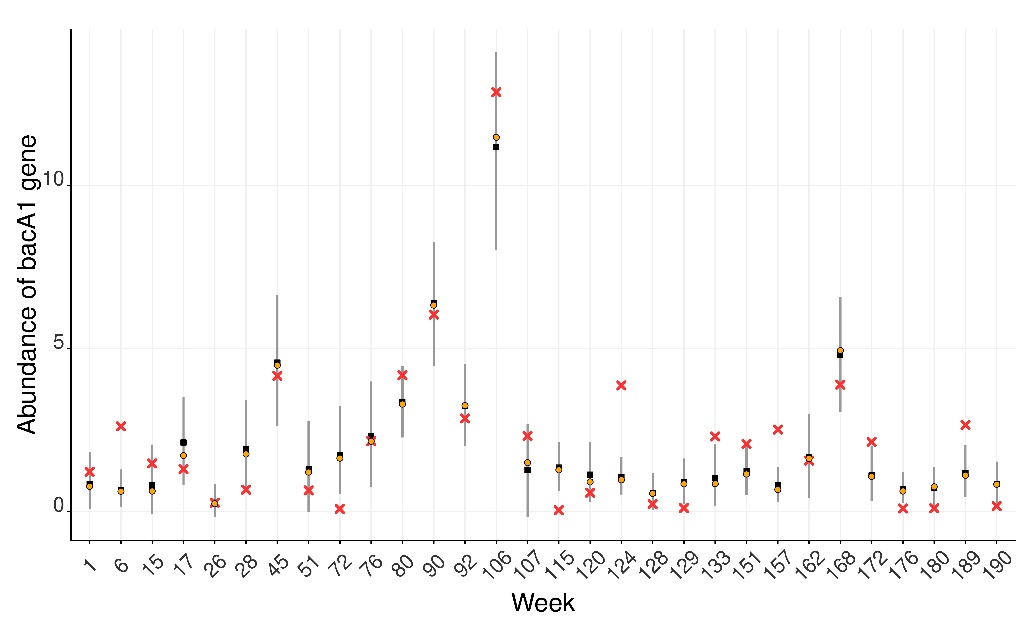

(c)

(b)

(a)

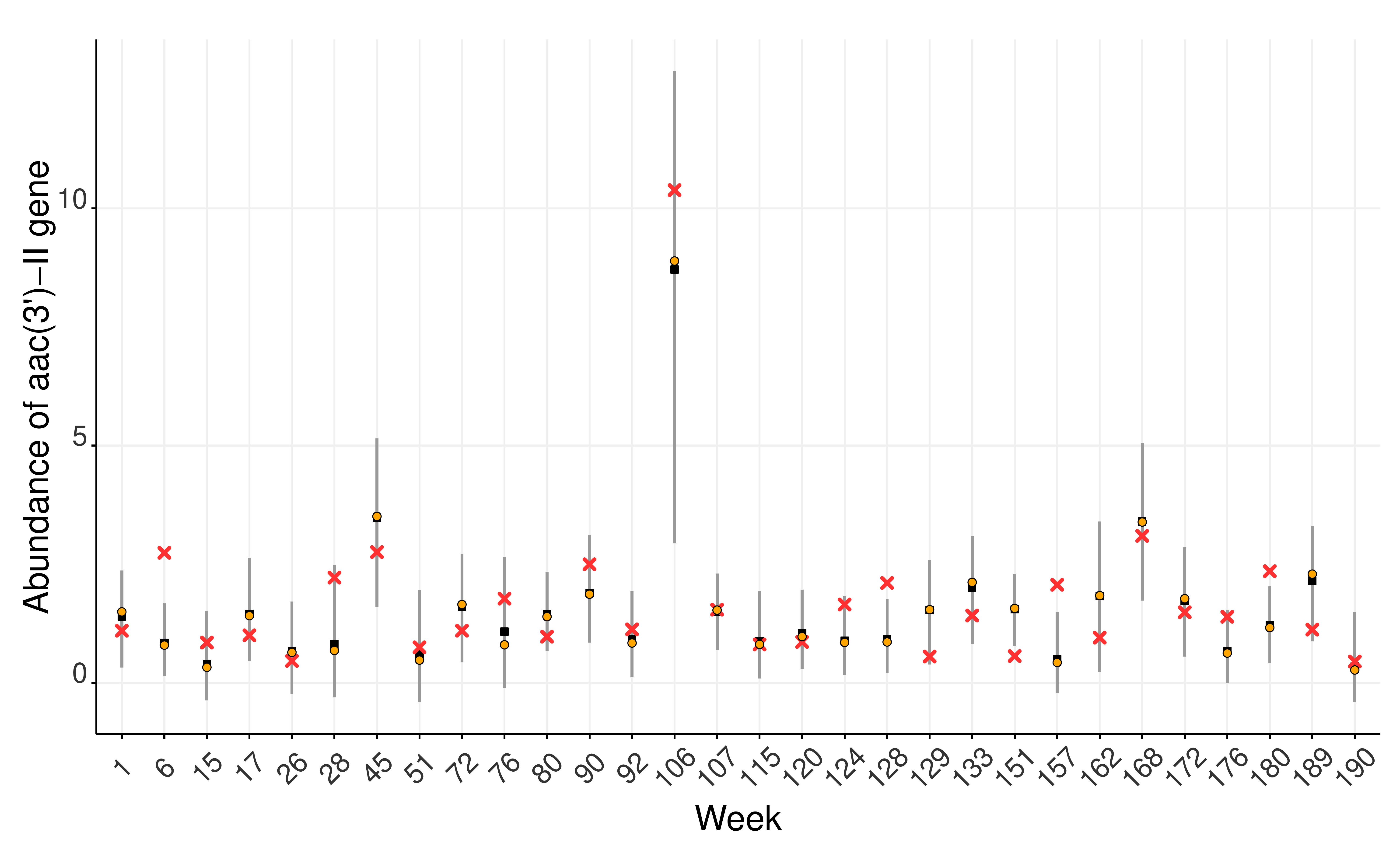

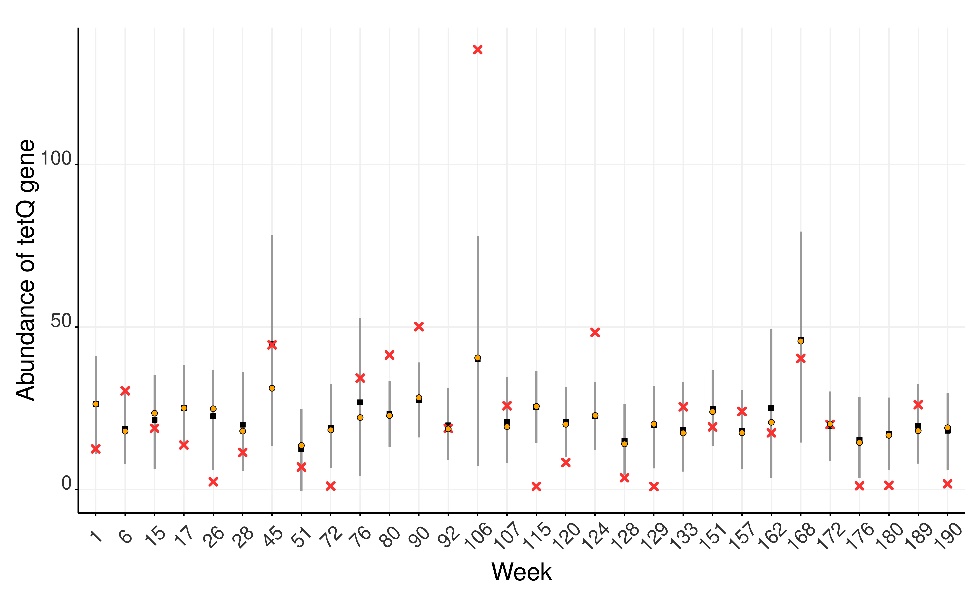

**Figure A.1.** Fitted models for gene abundance in untreated urban wastewater. (a) aac(3’)-II; (b) bacA1; (c) tetQ. Red crosses correspond to data, vertical grey lines to prediction intervals, black boxes to the median of output distributions, and orange circles to *K* (carrying-capacity) modes.

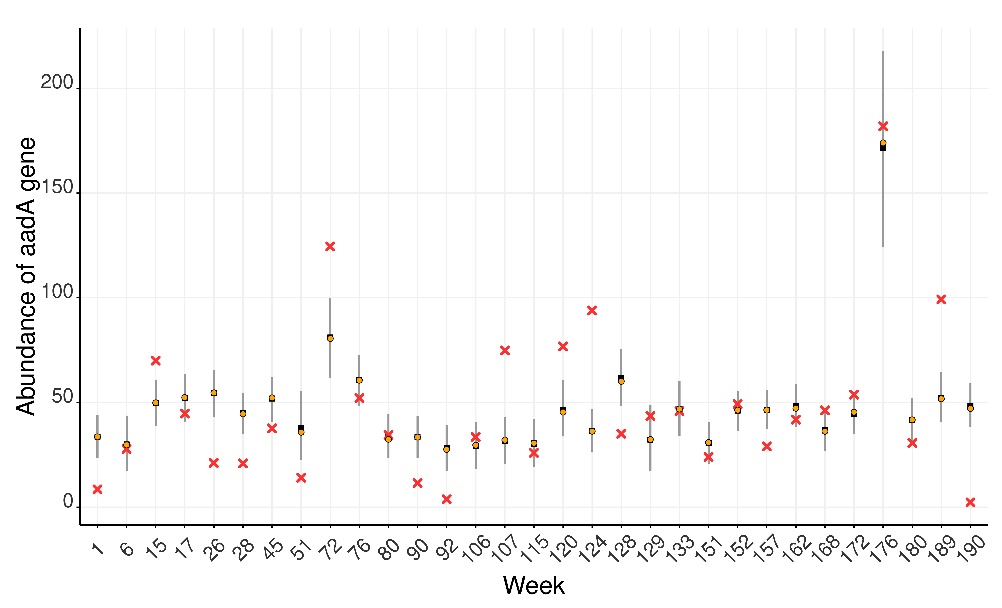

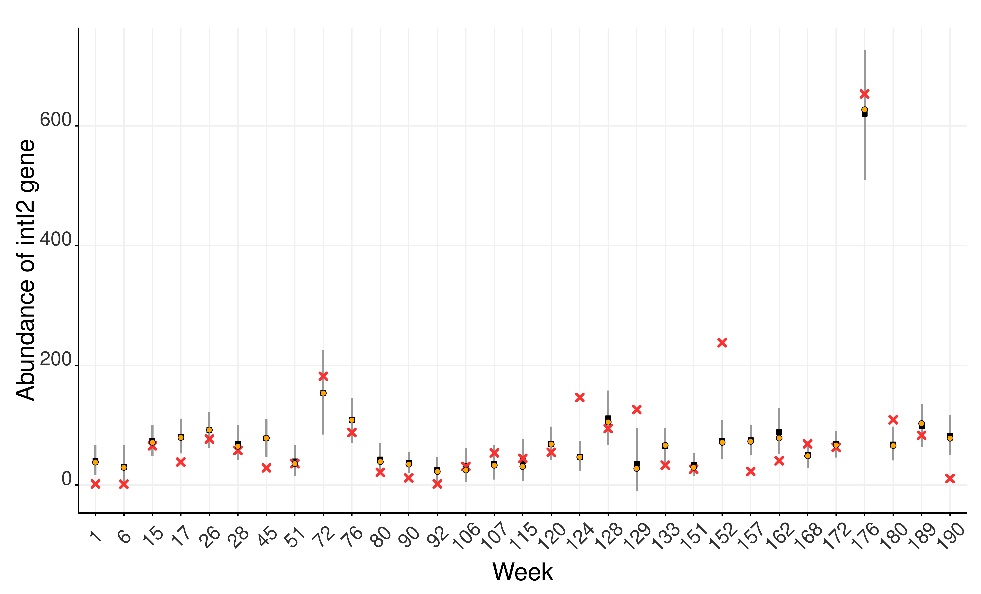

(c)

(b)

(a)

(d)

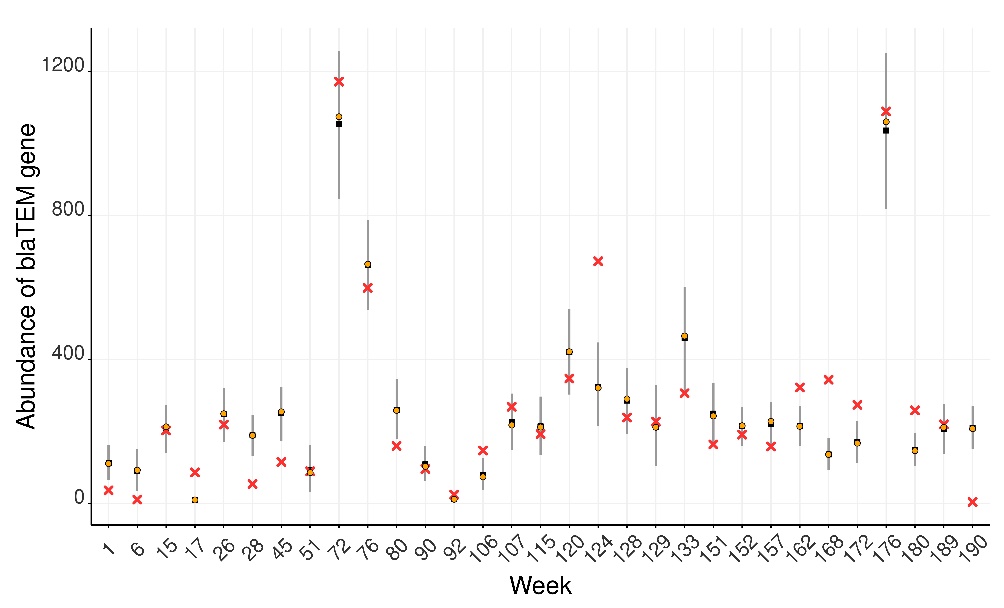

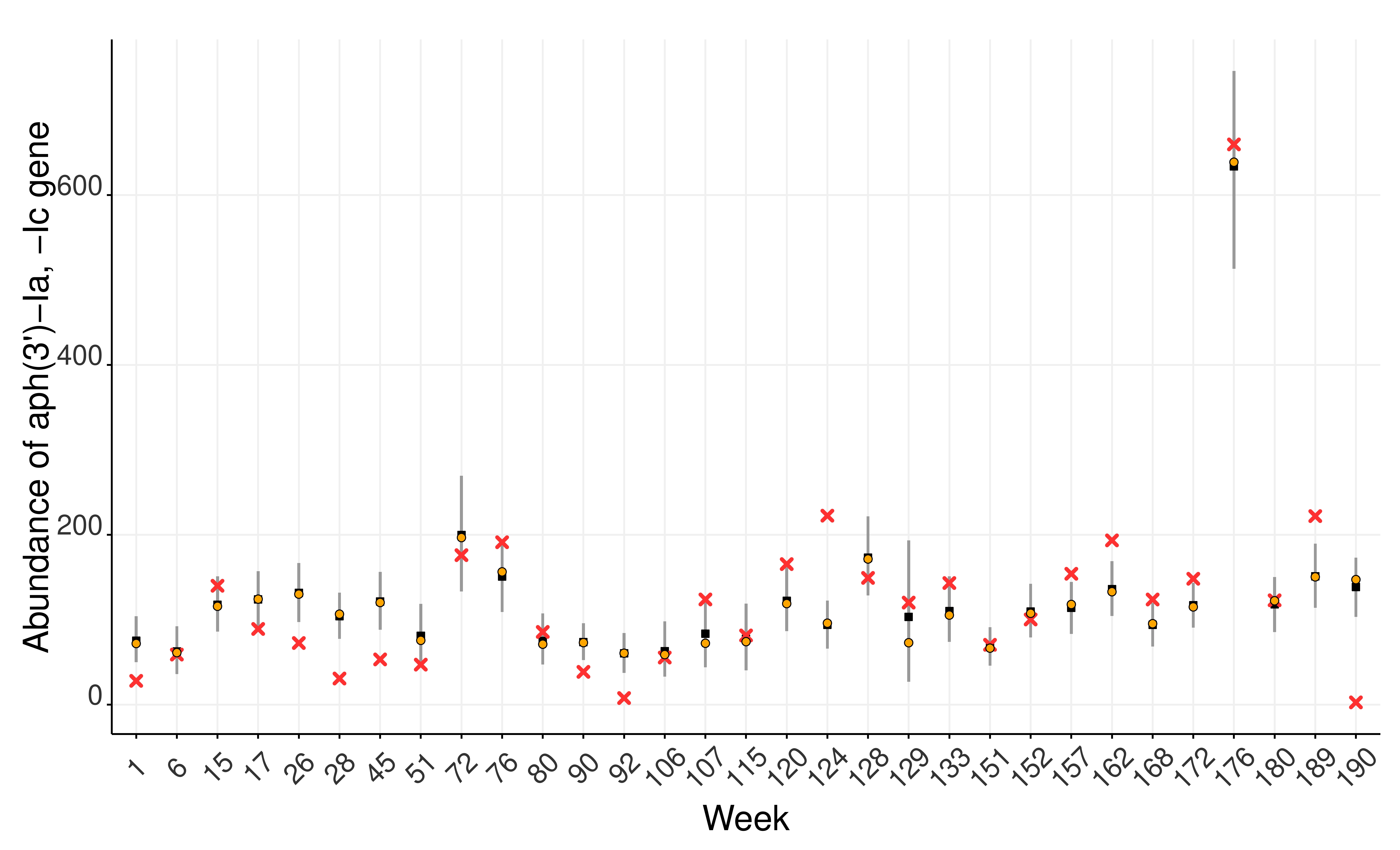

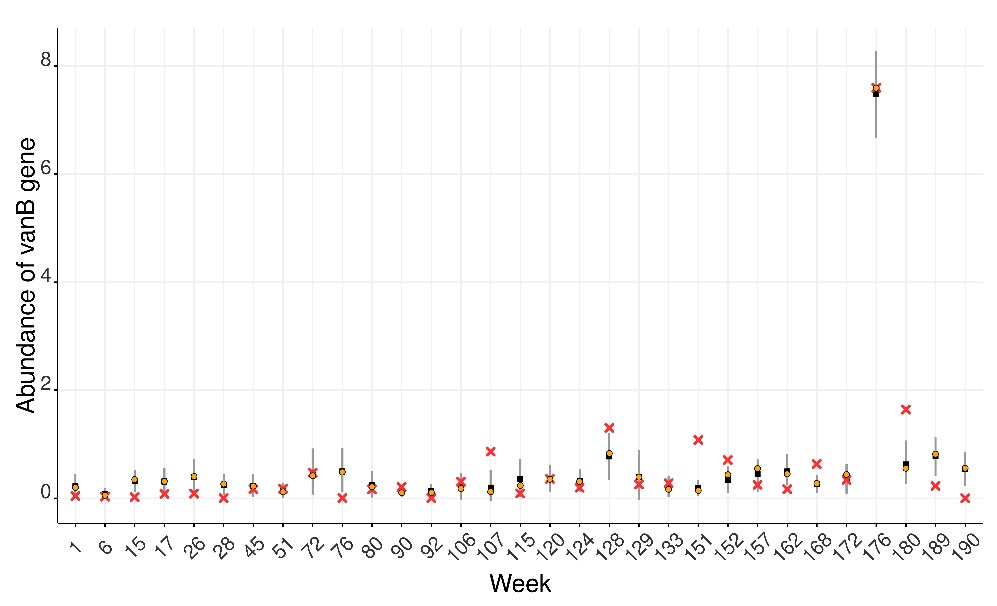

(e)

**Figure A.2.** Fitted models for gene abundance in untreated hospital wastewater. (a) aadA; (b) intI2; (c) blaTEM; (d) aph(3’) –la –lc**;** (e) vanB**.** Red crosses correspond to data, vertical grey lines to prediction intervals, black boxes to the median of output distributions, and orange circles to K (carrying-capacity) modes.

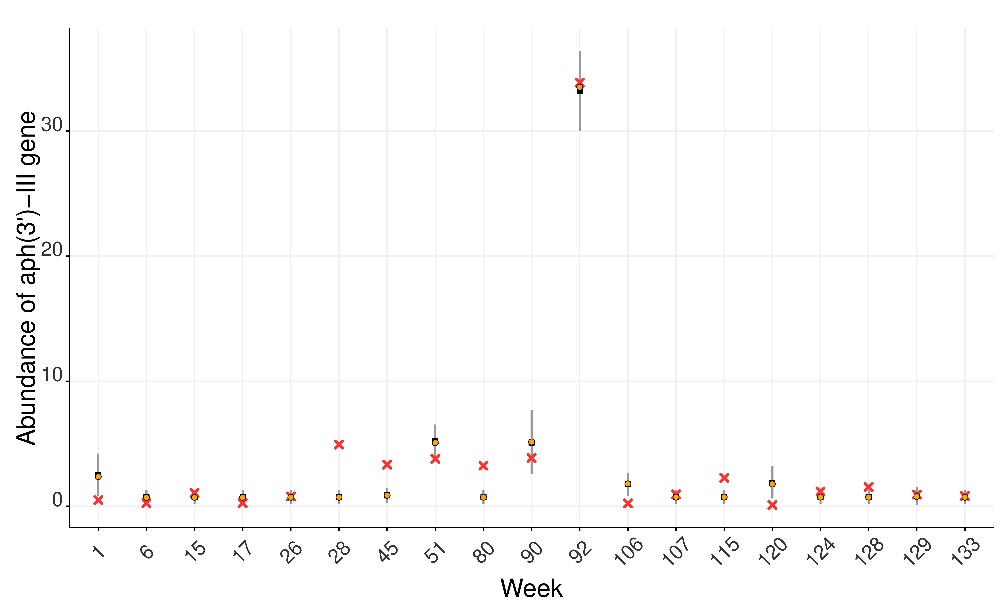

(b)

(a)

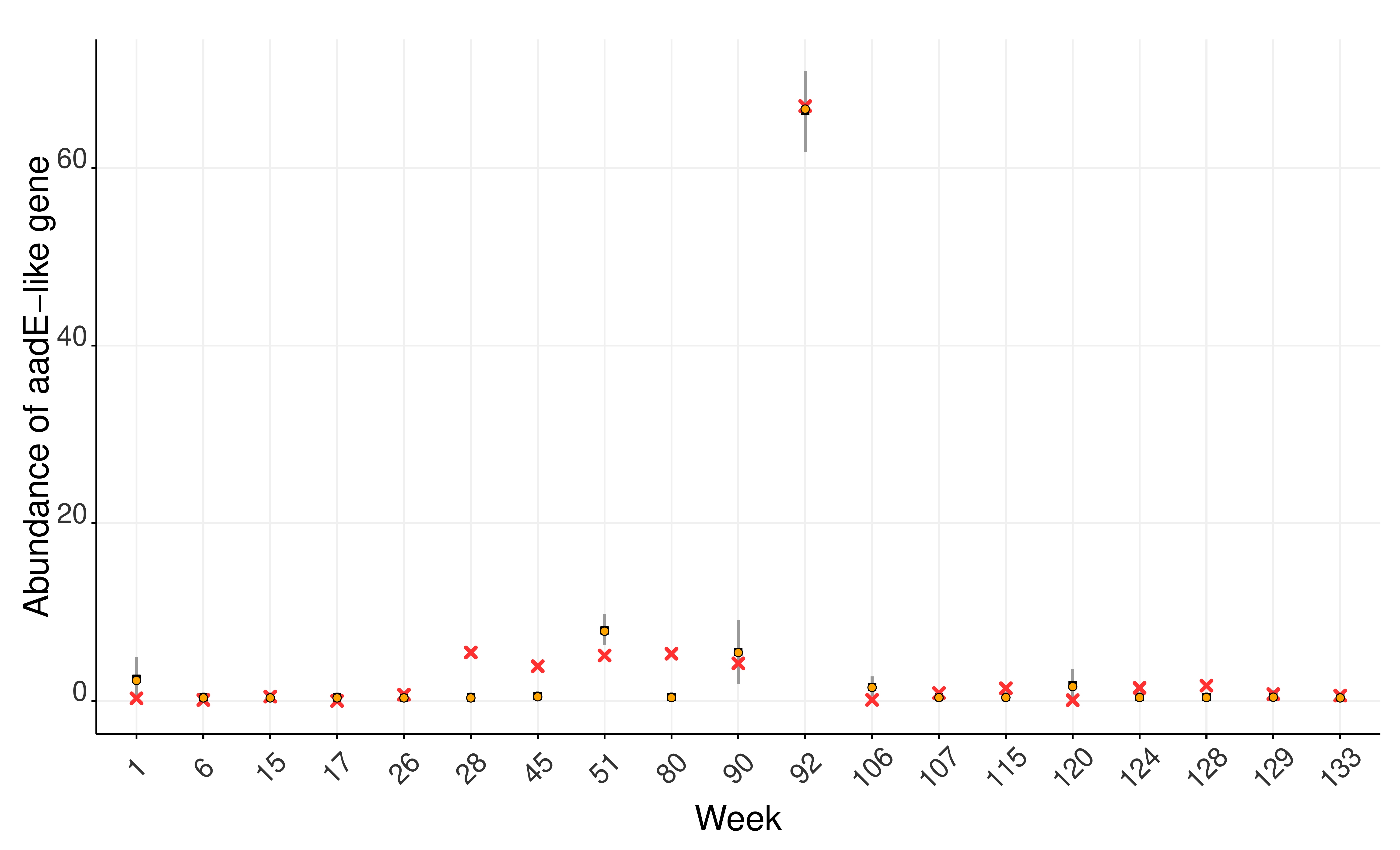

(d)

(c)

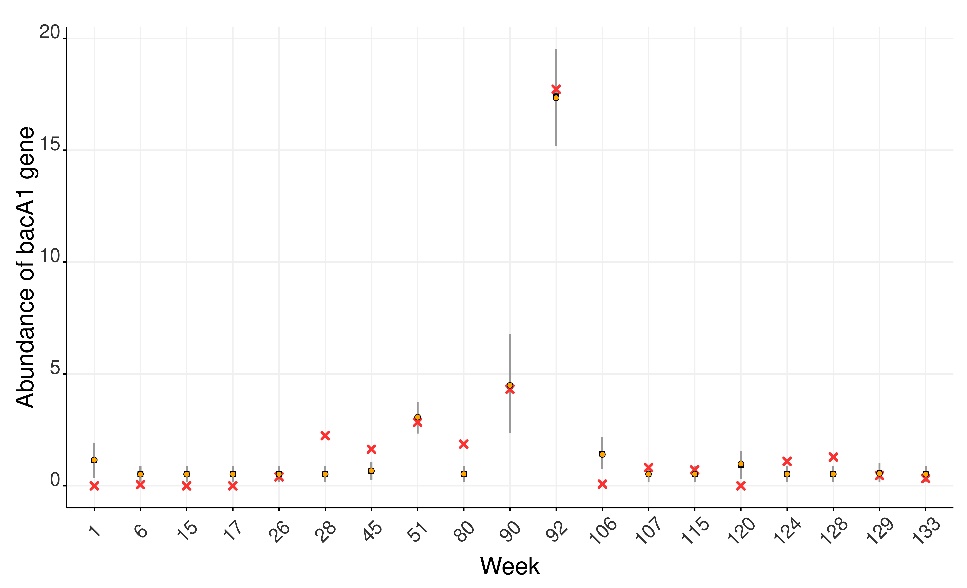

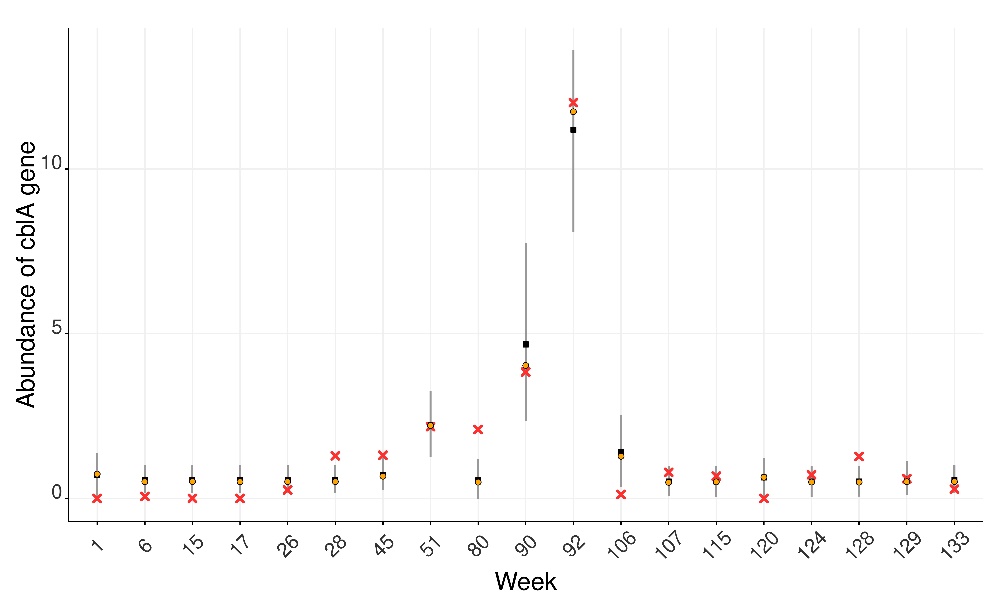

(f)

(e)
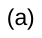
)

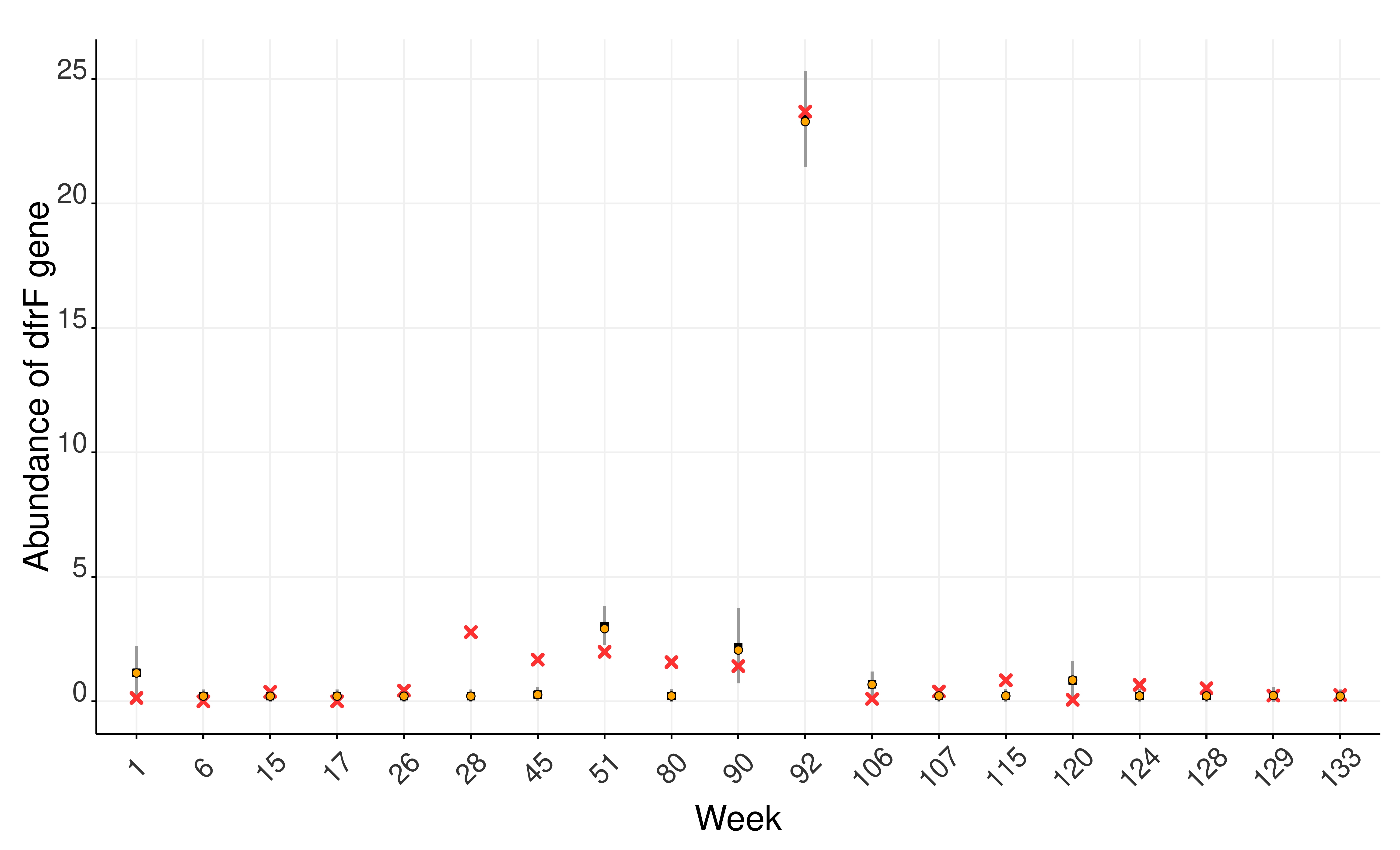

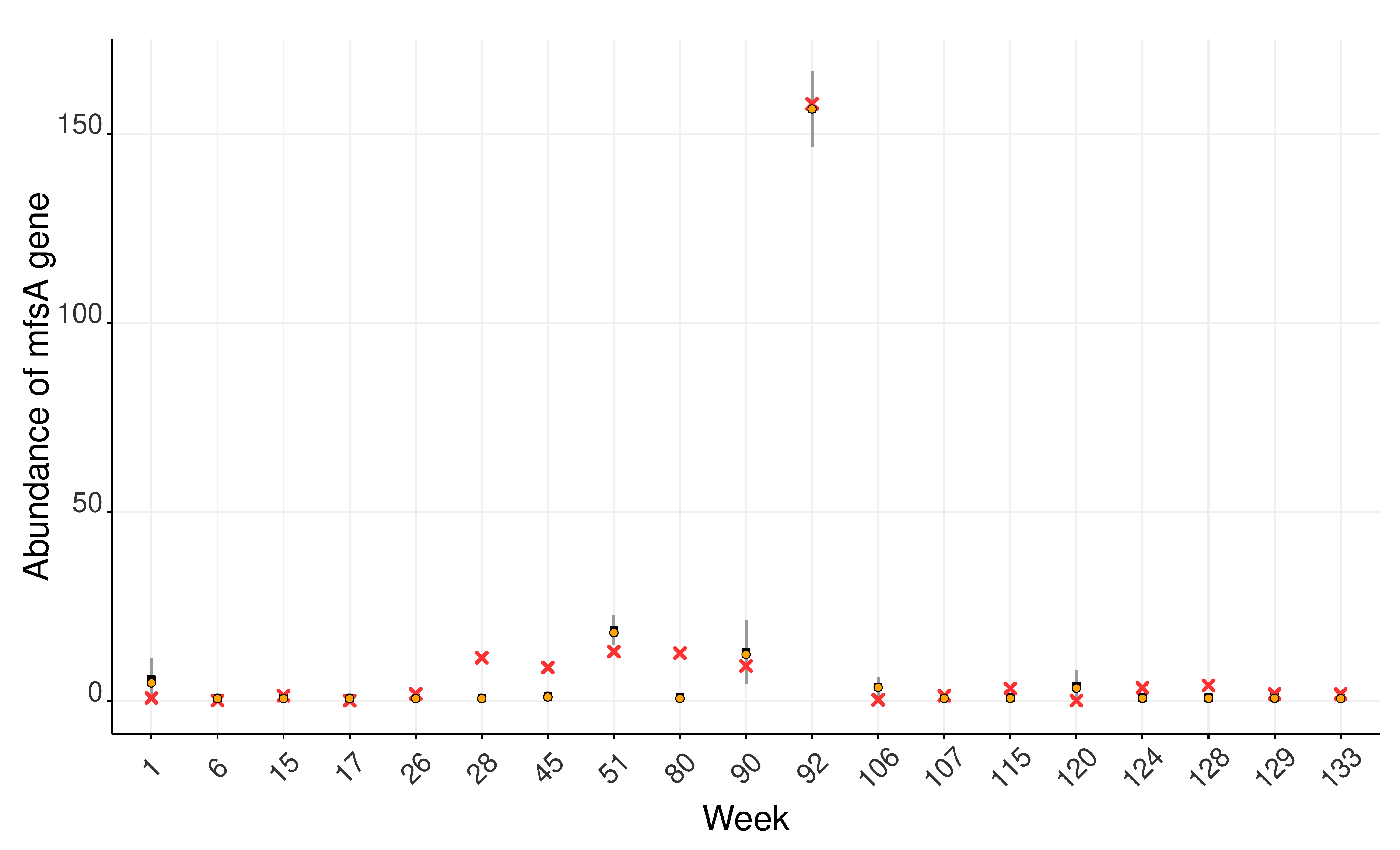

(h)

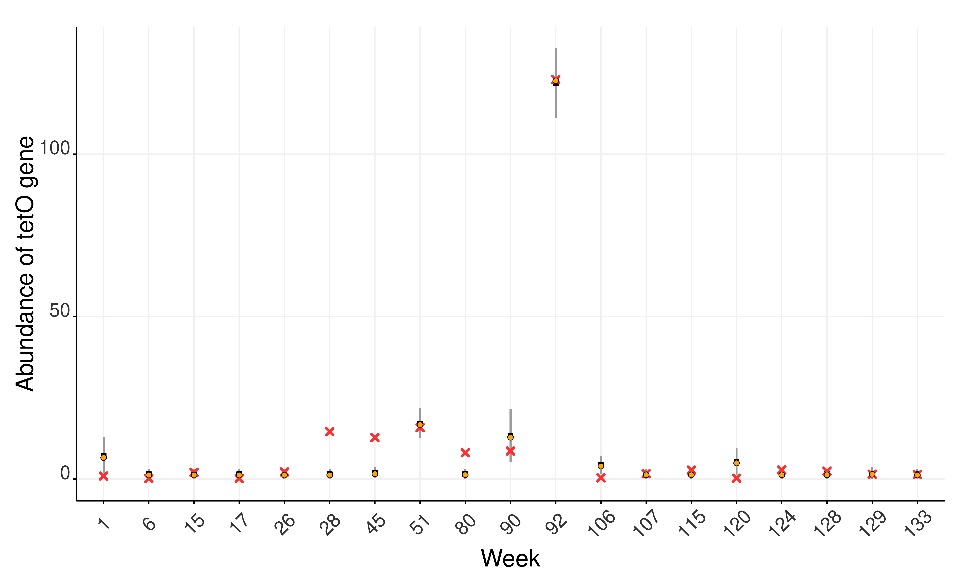

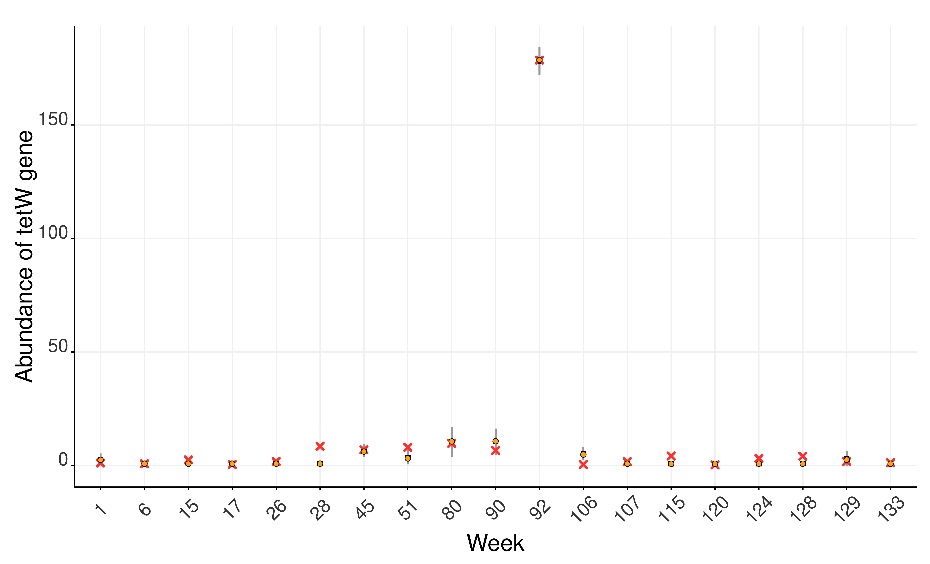

(g)

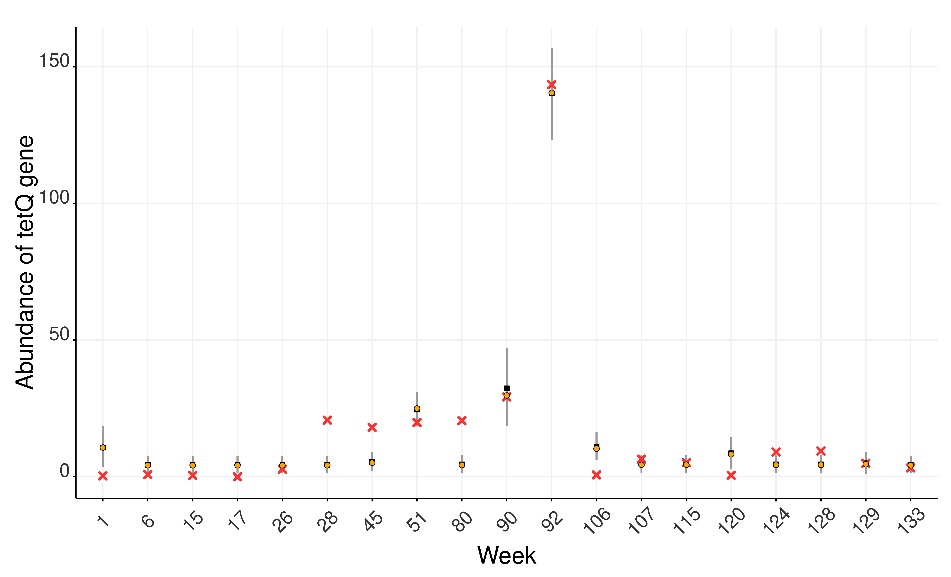

(i)

**Figure A.3.** Fitted models for the gene abundance in treated hospital wastewater. *(a) aadE-like; (b) aph(3’)-III; (c) bacA1; (d) cblA****;*** *(e) dfrF*; (f) *mfsA;* (g) *tetO*; (h) *tetQ*; (i) *tetW****.*** Red crosses correspond to data, vertical grey lines to prediction intervals, black boxes to the median of output distributions, and orange circles to *K* (carrying-capacity) modes.
